# Bacterial community associated with the dry-rot fungus *Serpula lacrymans* is dominated by Firmicutes and Proteobacteria

**DOI:** 10.1101/2020.11.24.397216

**Authors:** Julia Embacher, Sigrid Neuhauser, Susanne Zeilinger, Martin Kirchmair

## Abstract

The dry-rot fungus *Serpula lacrymans* causes enormous structural damage by decaying construction timber thereby resulting in tremendous financial loss. Dry-rot fungi decompose cellulose and hemicellulose and, if the wood remains wet, are often accompanied by a succession of bacteria and other fungi. Bacterial-fungal interactions have considerable impact on all interaction partners ranging from antagonistic to beneficial relationships. However, little is known about possible interaction partners of *S. lacrymans*. Here we show that *S. lacrymans* has many co-existing, mainly Gram-positive bacteria. By investigating differences in the bacterial community associated with fruiting bodies, mycelia and rhizomorphs, we provide evidence of preferential colonization of *S. lacrymans* tissues by certain bacterial phyla. Bacteria isolated from fruiting bodies and mycelia were dominated by Firmicutes, while bacteria isolated from rhizomorphs were dominated by Proteobacteria. Actinobacteria and Bacteroidetes were found in lower abundances. *In situ* fluorescence hybridization (FISH) analysis revealed that bacteria were not present biofilm-like, but occurred as independent cells, sometimes also attached to fungal spores. In co-culture, single bacterial isolates caused growth inhibition of *S. lacrymans* and vice versa. Additionally, certain bacteria induced pigment production in the fungus. Our results provide first insights for a better understanding of the holobiont *S. lacrymans* and give hints that bacteria are able to influence the behavior, e.g. growth and secondary metabolite production, of *S. lacrymans* in culture.

**Importance:** *Serpula lacrymans* is a very effective dry-rot causing fungus, specialized in degradation of coniferous timber in houses. The initial colonization is favored by water damage, and after establishment, the fungus starts to destruct cellulose and hemicellulose. It is among the most feared wood-rotting fungi in the built environment as the remediation of *S. lacrymans* damaged buildings is expensive and tedious. After improper renovation, the possibility of a recolonization by *S. lacrymans* is likely. As bacteria influence fungal establishment on wood, the need to investigate the bacterial community associated with *S. lacrymans* is apparent. The significance of our research is in identifying and characterizing bacteria associated with *S. lacrymans.* This will allow the assessment of their influence on fungal life style, leading to a broader understanding of the properties that make *S. lacrymans* so extraordinarily aggressive at decaying wood compared to other indoor wood destroyers.

## Introduction

The dry-rot fungus *Serpula lacrymans* is among the most feared wood decaying fungi in the built environment because of its ability to rapidly decompose cellulose and hemicellulose from coniferous wood and timber. Developmental stages of *S. lacrymans* include fruiting bodies (basidiomata – Fig. 1A), aerial mycelia (Fig. 1B) and rhizomorphs (Fig.1C). Fibre hyphae and generative hyphae (Fig. 1D, E) as well as ‘vessel’ hyphae (Fig. 1F) are observable in tissues with microscopy. The colonization of wood by *S. lacrymans* is characterized by fast growth of vegetative mycelia and the formation of thick (up to 2 cm diameter, Fig. 1C) mycelial cords that are used to transport nutrients and water to hyphae exploring new wood substrates (1). This allows *S. lacrymans* a rapid growth and makes it a successful invader in built environments (1, 3) resulting in particular economic importance of dry rot.

**Figure 1.**
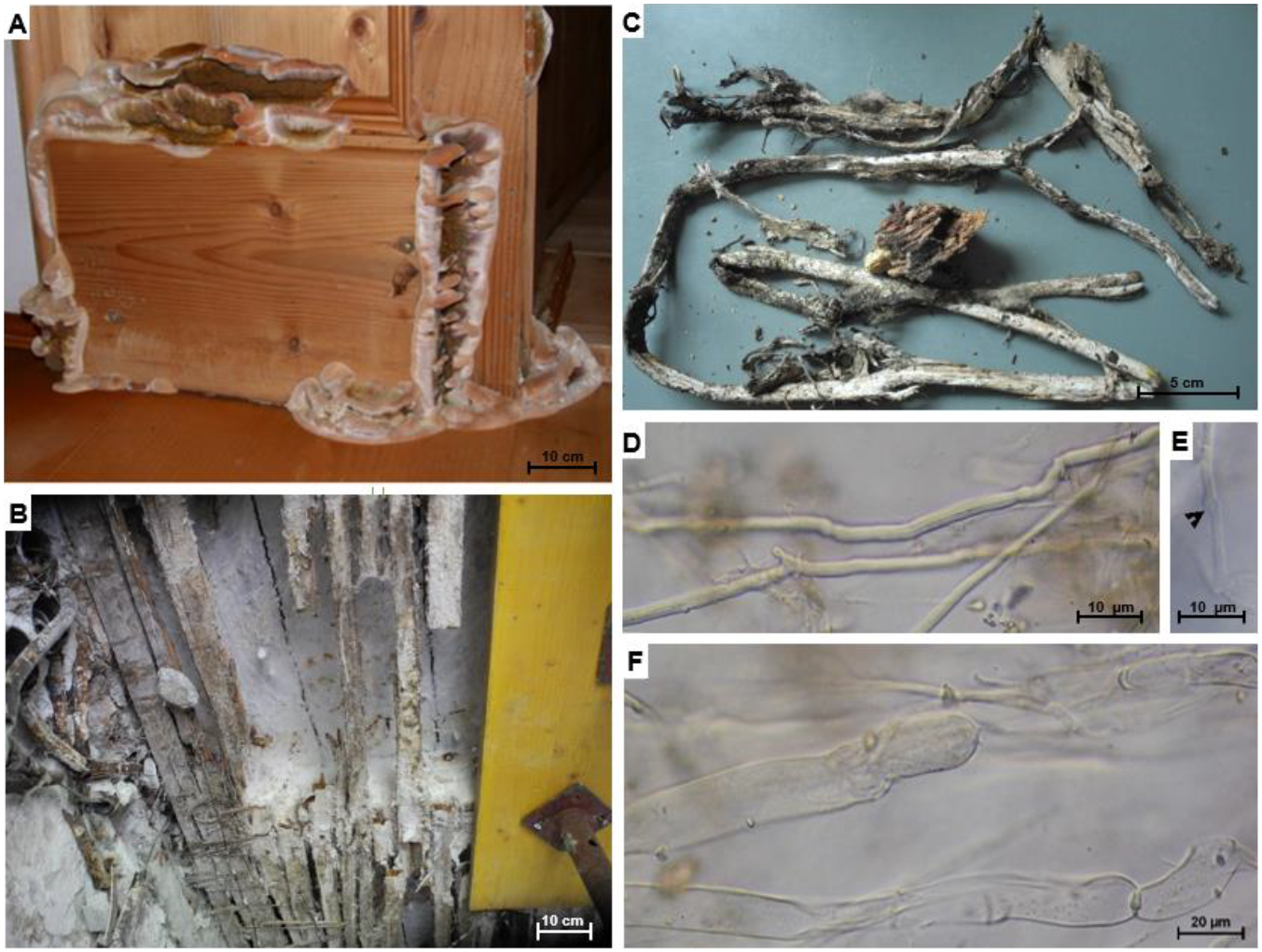
Infestation of timber by the dry rot fungus *Serpula lacrymans*. **A:** damaged wood with fruiting body **B:** fungal mycelia on wood beam ceiling.**C:** cord mycelium with wood **D:** fibre hyphae **E:** generative hyphae (arrow shows clamp-structures) **F:**‘vessel’ hyphae

Dry-rot fungi colonize wood/timber in buildings especially after water damage (4). Their establishment is influenced by bacteria, which are able to alter wood permeability and structure by production of e.g. cellulases and pectinases which open the crystalline structure of cellulose microfibrils (5) thereby increasing the susceptibility of wood for decay by fungi. Greaves (1971) was the first to develop a functional classification of wood-inhabiting bacteria: (i) bacteria that affect permeability but do not decrease material strength, (ii) bacteria that alter wood structures, (iii) bacteria that stimulate fungal decomposition; and (iv) bacteria that inhibit fungal decomposition (6). Colonization of wet wood by bacteria generally takes place at the beginning of the decay process (7, 8), although their establishment could be restricted by bactericidal components of ligninolytic fungi (9). Bacteria penetrate dead wood or timber via xylem parenchyma cells of ray tissues, vessels, tracheids and other wood cells, mainly using pits in cell walls for their penetration. There they can contribute to the degrading progress by living on pectin, monosaccharides and other easily accessible nutrients (10).

Nonetheless, fungi are considered as superior wood decomposers because of their larger size and mobility (via hyphal growth), and their marked capability to produce enzymes for degradation of the mayor building blocks of wood (lignin, cellulose, and hemicellulose) (7, 11). It was shown that wood decomposing bacteria live in close interaction with fungi (7, 12, 13), and that those bacteria are able to process low molecular mass sugars and small aromatic compounds that are released by ligninocellulolytic fungi (9, 11). There might be a synergistic interaction of bacteria and soft-rot fungi to predispose wood to *S. lacrymans* attack (7). It was discussed that the cellulase producer *Trichoderma viride* plays a role in timber degradation together with basidiomycetous wood-rotting fungi (14, 15). Timber degradation hence is likely a process to which different organisms are participating inter-, co- and counteracting with each other.

During wood degradation by basidiomycetes, environmental conditions become very challenging for bacteria because of rapid and strong acidification, production of reactive oxygen species, and toxic fungal secondary metabolites (16). In *Picea abies* logs, fungal diversity correlated negatively with bacterial abundance, while certain bacterial taxa co-occurred with certain fungi (17). Colonization of beech woodblocks with white-rot fungi *Hypholoma fasciculare* and *Resinicium bicolor* resulted in a strong bactericidal effect (18) while in soil basidiomycetes appeared to affect the community composition of bacteria (19, 20). Basidiomycete-associated bacteria need to survive these harsh conditions (16); hence specific soil bacteria must have evolved strategies to utilize fungal-secreted metabolites and overcome fungal defense mechanisms (21).

Bacterial-fungal interactions (BFI) can have different levels of specificity and a diverse range of interactions from antagonistic to beneficial relationships (Fig. S1). This means that the co-occurrence of bacteria and fungi is a result of physiological and metabolic interactions during which bacteria and fungi co-evolve and interdependently evolve. However, co-occurrence may be accidental and not representative of any causal relationship but the result of stochastic ‘mixing’. Any interaction between fungi and bacteria can modulate the behavior of neither, one or both of the interaction partners (22). Such BFI include competition for substrates (23) or the production of growth factors for fungi (7). The carbon-to-nitrogen ratio in wood is low and an increase in nitrogen by nitrogen-fixing bacteria living at the cost of carbohydrates set free by the fungi may be important (24, 25). Hence, the suite of bacteria that surround and interact with a fungus effectively constitutes its microbiome, and as such, they must be considered together (2). Therefore, the term holobiont is useful, as it is defined as “unit of biological organization composed of several distinct genomes, that, in principle, influence the genomic evolution of each other” (22, 26).

Only little is known about the natural bacterial and fungal interaction partners of *S. lacrymans*, although there are numerous fungal microbiome studies available (27-31). So far the analysis of airborne fungi in *S. lacrymans*-damaged homes revealed the co-occurrence of the ligninolytic *Donkioporia expansa* and the mycoparasite *T. viride* (14). Otherwise, many studies indicated that communication between interacting partners is key for any interaction. Communication molecules for bacteria and fungi include antibiotics and other secondary metabolites (SMs) produced by both partners which may be involved in mutualism, chemical warfare and in signaling (32). Fungus-associated bacteria have been shown to affect secondary metabolism of the fungus (33–37). A recent study analyzed the effect of bacteria on *S. lacrymans* pigment production (38, 39). Pigment synthesis from the quinone precursor atromentin, such as variegatic acid, was stimulated by 13 different bacteria and cell wall-lysing enzymes (e.g. β-glucanase, cellulase, proteases and chitinases), but not by lysozyme or mechanical damage. This may indicate a common pigmentation inducing mechanism, which could be triggered by fungal cell wall degradation. Moreover, the fungal pigments variegatic and xerocomic acid impact biofilm expansion and the swarming motility of bacteria such as *Bacillus subtilis (39).* During co-culturing with ubiquitous bacteria the gene cluster encoding for a synthetase and aminotransferase, both needed for atromentin biosynthesis, was induced (40).

The aim of this study was to investigate the abundance, composition and properties of the bacterial community of different *S. lacrymans* tissue types, including fruiting body, mycelia and rhizomorphs. We wanted to address (i) if the community of interaction partners is dominated by one or more phyla; (ii) if there is a tissue specific community composition; and (iii) if interaction partners change the behavior of *S. lacrymans* during co-culture experiments. The diversity of the bacterial community hosted by *S. lacrymans* was additionally investigated by *fluorescence in situ hybridization* (FISH) to get spatial information. The isolated bacteria were characterized regarding their ability for biopolymer degradation. We demonstrate that *S. lacrymans* has numerous bacterial partners, most of them Gram-positive, and provide first evidence that different tissue types are preferentially colonized by certain bacterial phyla. In co-culture assays, single bacterial isolates caused growth inhibition of *S. lacrymans* and vice versa.

## Results

### Microbial diversity associated with *S. lacrymans* fruiting bodies, mycelia and rhizomorphs

A total of 330 bacterial strains were isolated from 16 fruiting bodies, 9 rhizomorphs and 15 mycelium samples of *S. lacrymans*. The 16S rRNA sequence was generated for 301 of these bacterial isolates (Tab. S1). After removing sequences shorter than 400 bp, 274 sequences were aligned and their taxonomic affiliation and similarity was analyzed using phylogenetic analysis (Fig. S2-S7). Bacterial isolates belonged to 4 phyla, 8 classes, 9 orders, 27 families and 45 genera. Most 16S rDNA sequences had high similarity with type species (>98%-100%). The highest number of different species were within the Proteobacteria (22 species), followed by Actinobacteria (11 species) and Firmicutes (7 species). Bacterial diversity varied largely among the different tissues of *S. lacrymans*, with diversity being higher in fruiting bodies (27 species) and mycelia (25 species) compared to rhizomorphic tissue (14 species).

Results of the inverse morphotype approach showed that most bacterial isolates derived from *S. lacrymans* tissues were Gram-positive (1.2 x 10^10^ cells per g fresh weight; 58%). At the phylum level, Firmicutes (52% of all CFU) dominated the bacterial community. The second most abundant group were Proteobacteria (38%). Actinobacteria and Bacteroidetes were found to a lesser extent (6% and 4%, respectively). *Bacillus sp.* were most abundant (37%), followed by *Pseudomonas sp.* (16%) and *Kluyvera sp.* (13%). Bacteria isolated from fruiting bodies and mycelia were dominated by Firmicutes, while the rhizomorphic population was dominated by *Proteobacteria*. Rhizomorphs were predominantly colonized by *Pseudomonas sp.* (46%), while *Bacillus sp*. were most abundant (62%) on mycelia and *Kluyvera sp*. (30%) were the dominant species on fruiting bodies (Fig. 2).

**Figure 2.**
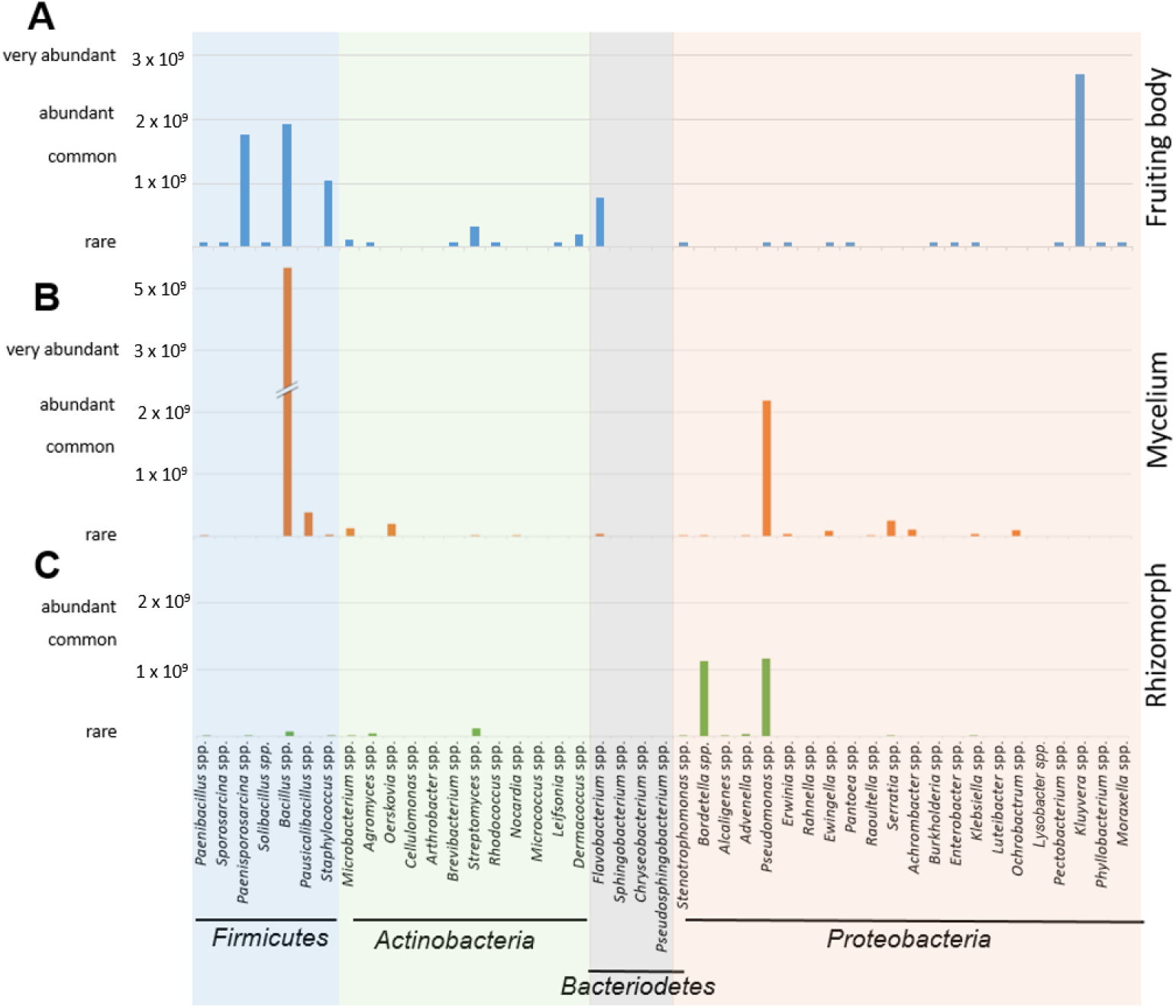
Gram-positive bacteria dominate the community on fruiting bodies, mycelia and rhizomorphs of *S. lacrymans*. Estimated abundances of bacterial species derived from different tissues of *S. lacrymans.* Because of the reciprocal approach only relative abundances are indicated. Based on estimated CFU counts bacteria were grouped as rare, common, abundant, and very abundant. Rare = 0 – 1×10^9^ CFU g^−1^ fungal tissue, common 1×10^9^ - 2×10^9^ CFU g^−1^, abundant = 2×10^9^ - 3×10^9^ CFU g^−1^ and very abundant = more than 3×10^9^ CFU g^−1^. Missing data means that there were either no identifiable colonies, because of too much similarity with other morphotypes, or no growth on agar plates. Abundances of bacterial species (in CFU g^−1)^ were estimated by using an inverse morphotype approach. **A**: isolates from fruiting bodies **B:** mycelia **C:** rhizomorphs

The CFU g^−1^ counts on tryptone soy agar (TSA) and R2A agar were comparable. The total amount of CFU g^−1^ varied in the different tissue types, with the highest CFU counts derived from mycelium (Fig. 3A). On the other hand, the variation was high between sampling sites (Tab. 1: no. 8, 9, 10, 11, 15 and 18 vs. 12, 13, 16 and 17) (Fig. 3B).

**Figure 3.**
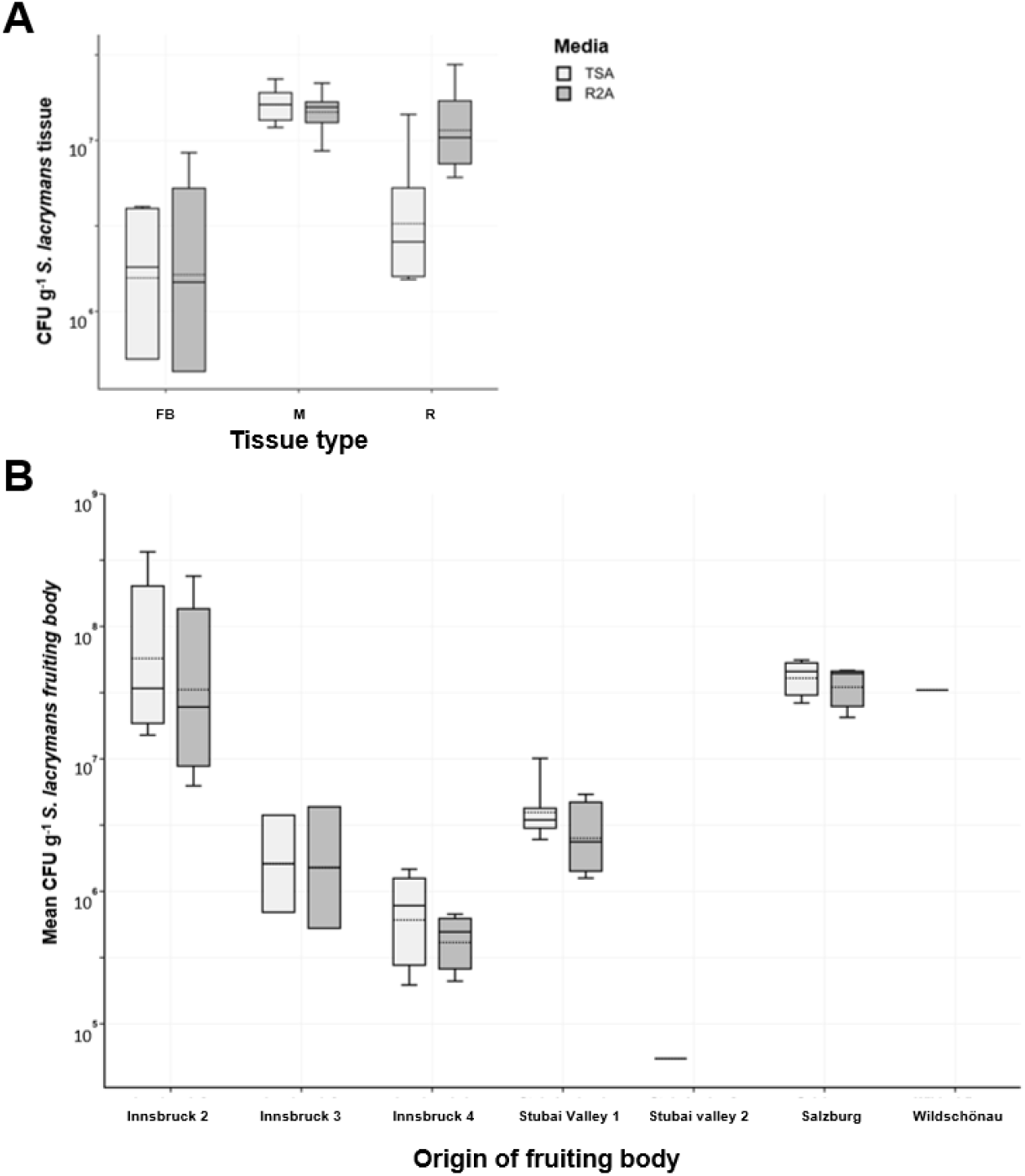
The number of bacterial cells on *S. lacrymans* tissue depends on tissue type and sampling site. **A** Absolute abundance of bacterial cells (in CFU g^−1^) derived from different tissues of *S. lacrymans* Innsbruck 3 (table 1, No. 12), namely fruiting bodies (FB), mycelia (M) and rhizomorphs (R). **B** Mean CFU g^−1^ of *S. lacrymans* fruiting bodies (Innsbruck 1 - 7, Stubai valley 2, Salzburg and Wildschönau). Data are displayed in logarithmic scale. Bacteria were cultured on tryptic soy agar (TSA, light grey columns) and R2A agar (R2A, dark columns).

**Table 1:** Description of collected material of *Serpula lacrymans* for bacterial isolation

| No. | Origin | Fruiting body | Mycelia | Rhizomorph | Date |
| --- | --- | --- | --- | --- | --- |
| 1 | Außerfern | 2 | 1 | 1 | 13 December 2018 |
| 2 | Brixen valley 1 | 0 | 2 | 0 | 19 October 2018 |
| 3 | Brixen valley 2 |  | 1 |  | 02 May 2019 |
| 4 | Brixen valley 2 | 1 | 1 |  | 19 June 2019 |
| 5 | Buchkirchen | 1 |  | 1 | 14 June 2019 |
| 6 | Grieskirchen | 1 | 1 | 1 | 13 June 2019 |
| 7 | Innsbruck 1 | 2 | 3 | 2 | 07 November 2018 |
| 8 | Innsbruck 2 | 2 |  |  | 22 March 2019 |
| 9 | Innsbruck 2 | 1 |  |  | 2 July 2019 |
| 10 | Innsbruck 2 | 1 | 1 | 1 | 16 August 2019 |
| 11 | Innsbruck 2 | 1 |  | 1 | 06 September 2019 |
| 12 | Innsbruck 3 | 1 | 1 | 1 | 30 October 2019 |
| 13 | Innsbruck 4 | 1 |  |  | 18 November 2019 |
| 14 | Pitz valley | 1 | 1 | 1 | 16 May 2019 |
| 15 | Salzburg | 1 |  |  | 31 October 2019 |
| 16 | Stubai valley 1 | 1 |  |  | 14 October 2019 |
| 17 | Stubai valley 2 |  | 1 | 1 | 28 November 2019 |
| 18 | Wildschönau | 1 | 0 | 0 | 27 June 2018 |
| 19 | Traunkirchen |  |  | 1 | 26 June 2020 |
| 20 | Zillertal valley | 1 | 1 |  | 2 July 2020 |

### Detection and localization of bacterial interaction partners with Fluorescence In Situ Hybridization (FISH)

Spherical bacteria (0.5 to 1.6 µm diameter) could be visualized on fruiting bodies, mycelium, and rhizomorphic structures of *S. lacrymans.* Bacteria were present mainly as individual cells (Fig. 4 and 5) and not as biofilms. 58% of all signals detected with the multiplexed FISH assay were derived from the LGC354ABCw-CY5-probe, indicating that the majority of all visualized bacteria belonged to the phylum Firmicutes. About 13% of all signals were visualized with the BET42aw-6FAM and GAM42aw-6FAM probes (β- and γ-Proteobacteria), and 29% of signals originated from the general Eubacterial probe (Fig. 6). The analysis of all analyzed slides (multiplex FISH) gave a mean of 24 signals per section. The medium surface area and thickness of the fungal sections were used to extrapolate and estimate the number of bacteria per cm^3^ fungal structure. Based on the approximation that 1 cm^3^ of fungal fruiting body equals 1 g, 24 signals per section add up to 6.6 x 10^7^ CFU g^−1^ fruiting body. These results are similar to the isolation-based approach, where 10^5^ to 10^8^ CFU g^−1^ fruiting body were identified (Fig. 3).

**Figure 4.**
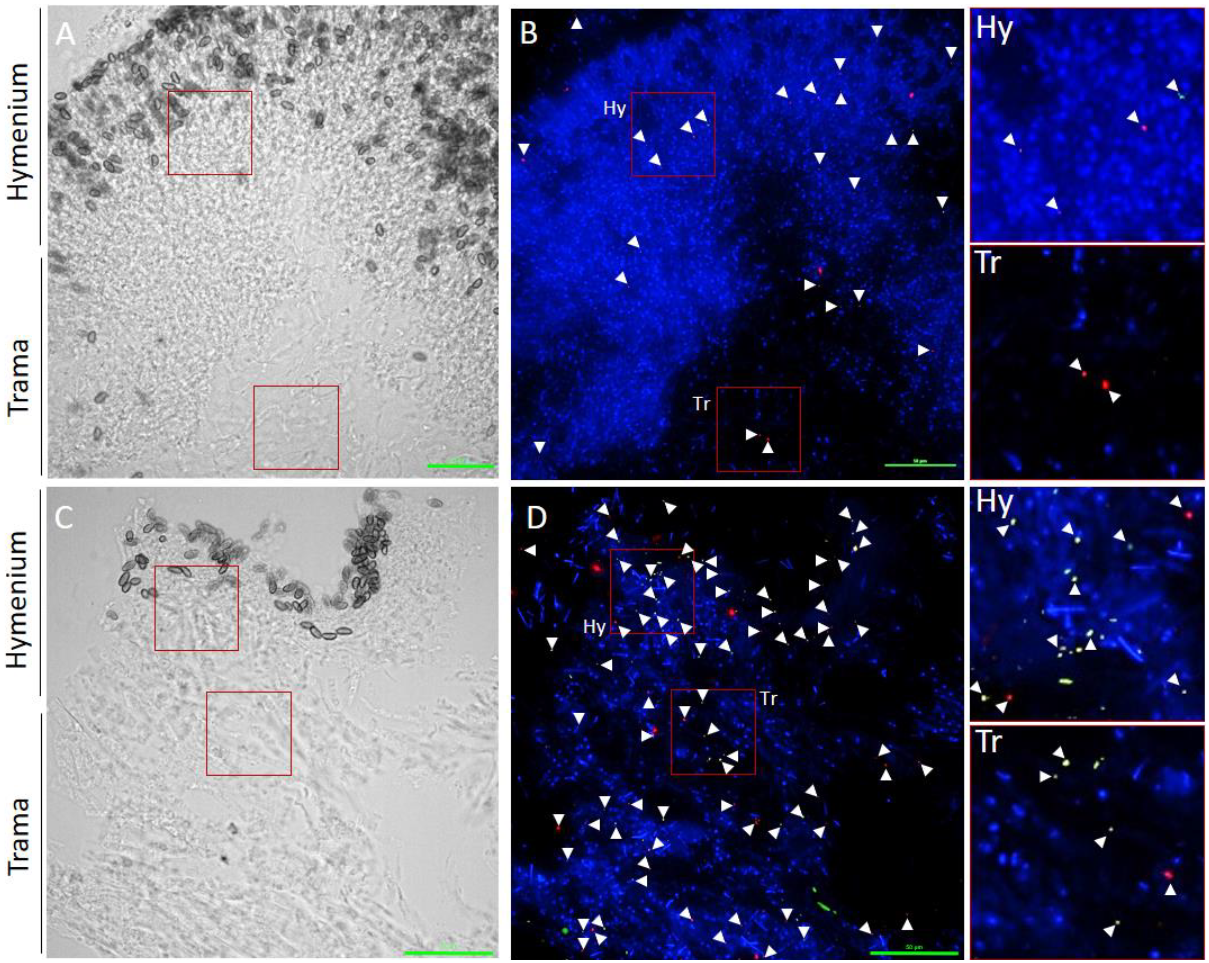
Localization of bacteria on *S. lacrymans* fruiting bodies. EUB338 Mix-Cy3, BET425aw-FAM, GAM42aw-FAM, and LGC354 Mix-Cy5 bacterial FISH probes were used. Firmicutes are displayed as red objects, β- and γ-Proteobacteria as green objects, and other bacterial cells in orange. **A** Transmitted light picture. **B** Overlay of DAPI channel and FISH probe channels. **C** Transmitted light picture. **D** Overlay of DAPI channel and FISH probe channels. Pictures on the right side are enlargements from B and D. Signals are highlighted with white arrows. Scale bars are 50 µm.

**Figure 5.**
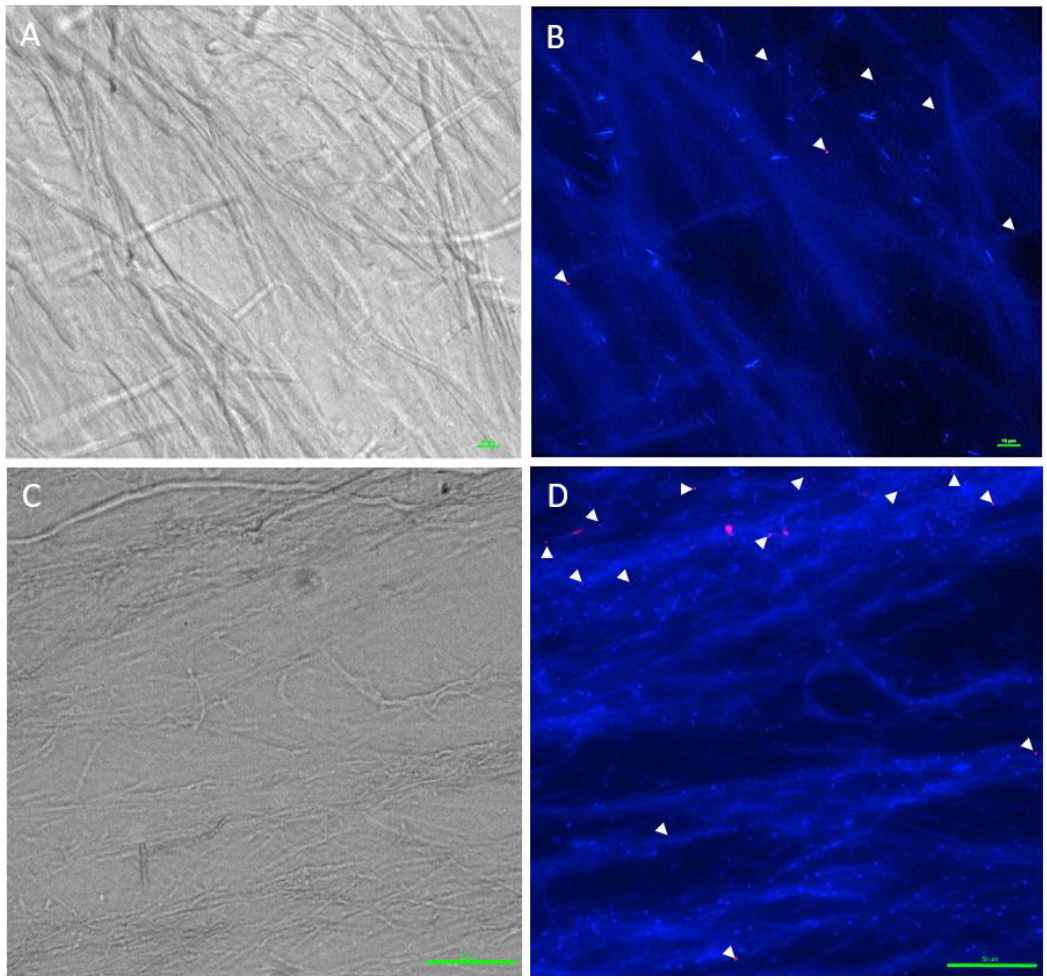
Localization of bacteria on *S. lacrymans* rhizomorphs and mycelium. **A** Overlay of signals derived from EUB338 Mix-Cy5 and DAPI. Investigated tissue type = rhizomorph. **B** Transmitted light picture of rhizomorph. **C** Overlay of signals derived from EUB338 Mix-Cy5 and DAPI. Investigated tissue type = mycelium. **D** Transmitted light picture of fungal mycelium. The general bacterial EUB338 Mix probe gave red signals. Signals are highlighted with white arrows. Scale bars are 10 and 50 µm.

**Figure 6.**
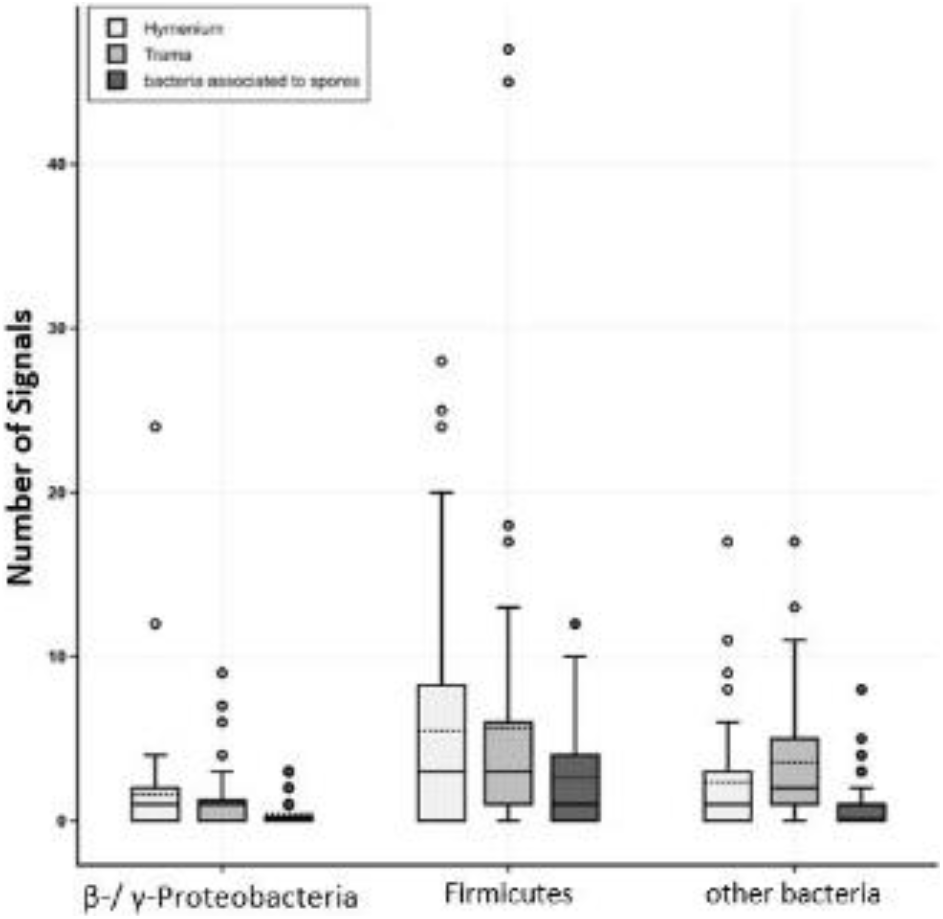
FISH data shows dominance of Firmicutes on *S. lacrymans* fruiting bodies. Probes were applied in a multiplexed way. Fungal fruiting bodies from Salzburg, Innsbruck 1 and 4 (Austria; table 1 no. 7, 13, and 15) were investigated in triplicates. Obtained signals were split into the categories hymenium or trama of fruiting bodies and bacteria associated with spores. Positive signals for β- and γ-Proteobacteria (BET42aw-6FAM and GAM42aw-6FAM probes), Firmicutes (probe LGC354ABCw-CY5) and other bacteria (probe EUB338 Mix-Cy3) are displayed.

The hymenium contained 45% of bacterial signals, and 34% of bacterial signals were located in the trama of the fruiting bodies. Approximately 17% of all visualized bacterial cells were associated with fungal spores. The fruiting body from Salzburg (Tab. 1, no. 15) was not entirely mature and therefore had no real hymenium, thus this fruiting body was excluded from these calculations. Notably, the number of detected signals was relatively balanced regarding the three tested fruiting bodies collected at different locations; 33% of all signals emerged from the Innsbruck 4 fruiting body (Tab. 1, no. 13), 36% from that collected in Salzburg (Tab. 1, no. 15), and 31% from the fruiting body from Innsbruck 1 (Tab. 1, no. 7). However, the immature fruiting body (Salzburg) showed a higher content of β- and γ-Proteobacteria (20% vs. 6% and 12% in the mature fruiting bodies). Investigation of mycelial and rhizomorphic tissues with different probes revealed no spatial pattern of bacterial dispersal (Fig. 5). Importantly, neither did we find bacteria embedded in a biofilm-like, continuous layer nor any evidence for intracellular bacteria.

### Enzymatic activities of isolated bacteria

Bacteria that were able to depolymerize pectin, xylan, starch or cellulose showed a clear halo-like zone around their colonies when cultivated on plates containing the respective biopolymer (Fig. 7). In total, 10 of the 327 tested bacteria showed activities of all four enzymes. These were *Serratia* sp. (No. 13), *Paenibacillus* spp. (No. 36 and 79), *Flavobacterium* spp. (No. 41 and 46), one *Arthrobacter* sp. (No. 69), one *Bacillus* sp. (No. 279) and two *Pseudomonas* spp. (91 and 221). Isolate No. 366 b was able to degrade all tested carbohydrates but could not be identified on the phylogenetic level. 25% of the isolates were able to degrade pectin, 43% xylan, 17% carboxymethylcellulose, and about 66% were able to depolymerize starch (for detailed information see table S2 in the supplement); thus the ability to degrade starch was found most commonly among the bacterial isolates.

**Figure 7.**
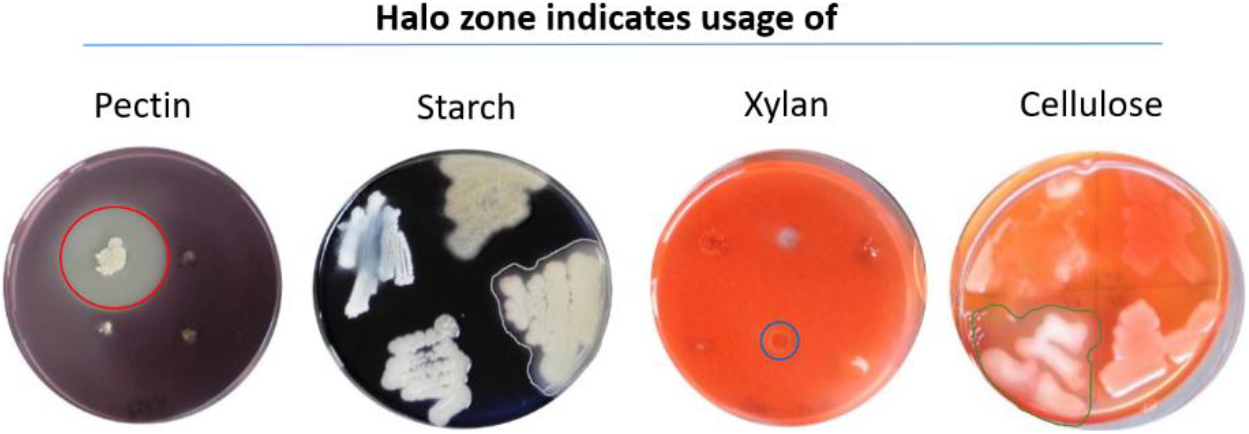
Production of carbohydrate degrading enzymes (pectinases, amylases, xylanases, or cellulases). Hydrolysis zones indicate that the bacteria can use xylan (blue circle), starch (grey border), pectin (red circle) or carboxymethyl cellulose (green border) as carbon source. Used media: pectinase screening agar medium (PSAM), starch degradation screening agar (SSA), xylanase screening agar (XSA) and carboxymethyl cellulose agar (CMC). Plates were incubated at 25 °C. Substrate degradation halos were visualized with 0.1% congo red solution or potassium iodide-iodine solution.

### Effects of co-cultivation on fungal and bacterial growth

After 31 days of co-incubation of *S. lacrymans* with each of the 50 bacterial isolates (Tab. 2), 16 interactions emerged as neutral or had a slightly positive effect on mycelial growth of the fungus, with growth rates of 90 – 117% compared to the mean growth of the control experiment (Fig. 8A). Four isolates were moderately growth-inhibiting (60 - 90% growth compared to the control, Fig. 8A) while 25 isolates strongly inhibited mycelial growth of *S. lacrymans* (less than 60% growth in comparison to the control, Fig. 8B). Bacteria which had the highest growth-inhibiting effect on *S. lacrymans* were *Microbacterium* sp. (isolate no. 1, 45, and 95), *Arthrobacter* sp. (isolate 69), *Bacillus* sp. (isolate no. 71 and 98), *Paenibacillus* sp. (isolate 36), *Oerskovia* sp. (isolate 106) and *Streptomyces* sp. (isolate 95) (Fig. 8B; last nine isolates). Among the five bacteria strongly inhibiting the growth of *S. lacrymans,* four belonged to *Bacillus* sp. (isolates 71, 87, 98 and 107), which were identified as the dominating group isolated from *S. lacrymans* structures (Fig. 2). One *Bacillus* strain (No. 54), however, was showing neutral behavior. The co-cultivations were again analyzed after 50 days, but no changes compared to the 31 days analyses emerged.

**Figure 8.**
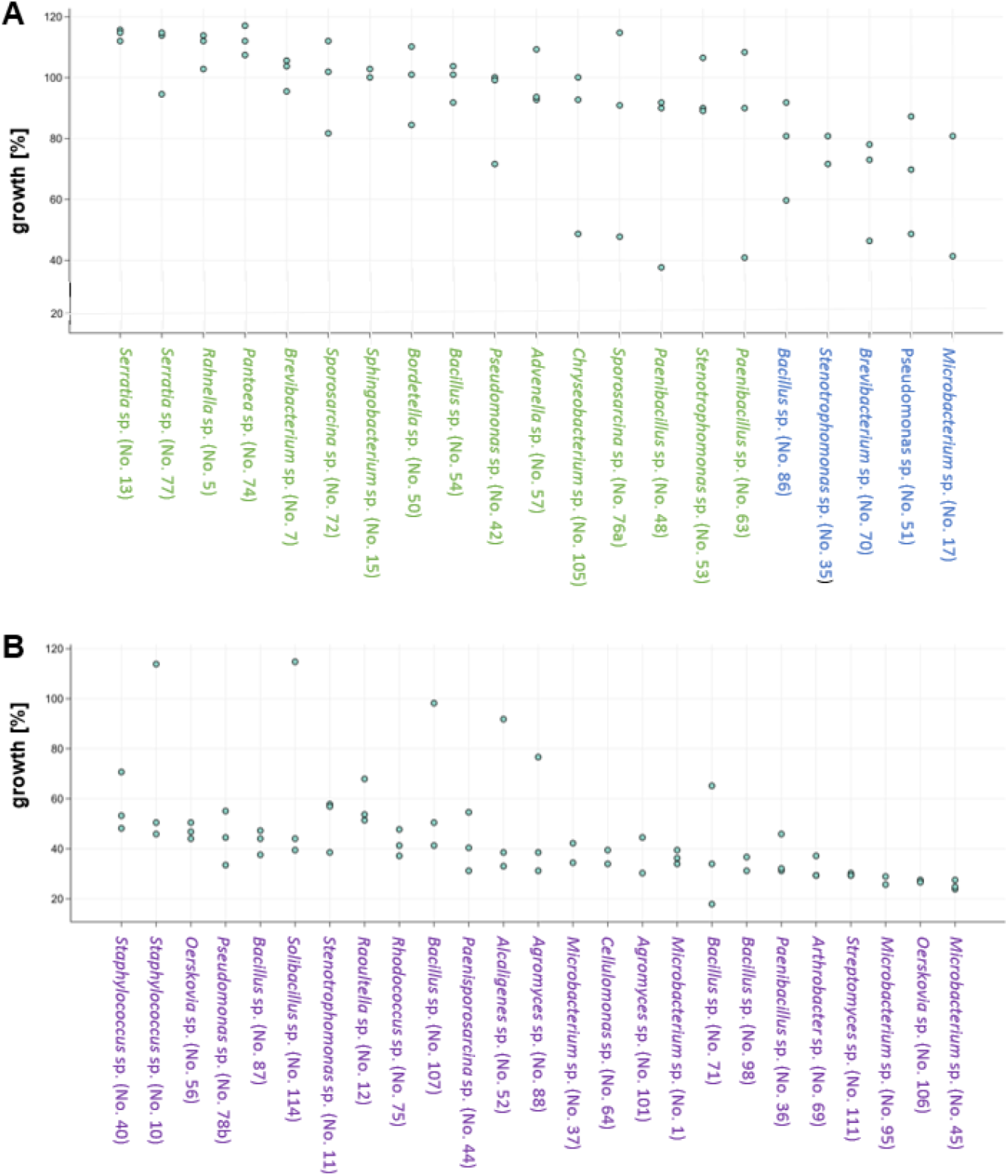
Bacterial co-cultivation partners can impair the growth of *S. lacrymans*. Fungal growth (in %; in comparison to the growth controls) upon co-incubation with selected bacteria on minimal media. Results from the analysis after 31 days of co-culture are displayed. Two to three replicates per bacterial isolate were performed. Bacteria from left to right are more growth inhibiting. **A** Isolates with neutral (90-120% growth; green font) or inhibiting (60-90%; blue font) effects. **B** Isolates that were categorized as strongly inhibiting fungal growth (growth less than 60%; purple font).

**Table 2:**
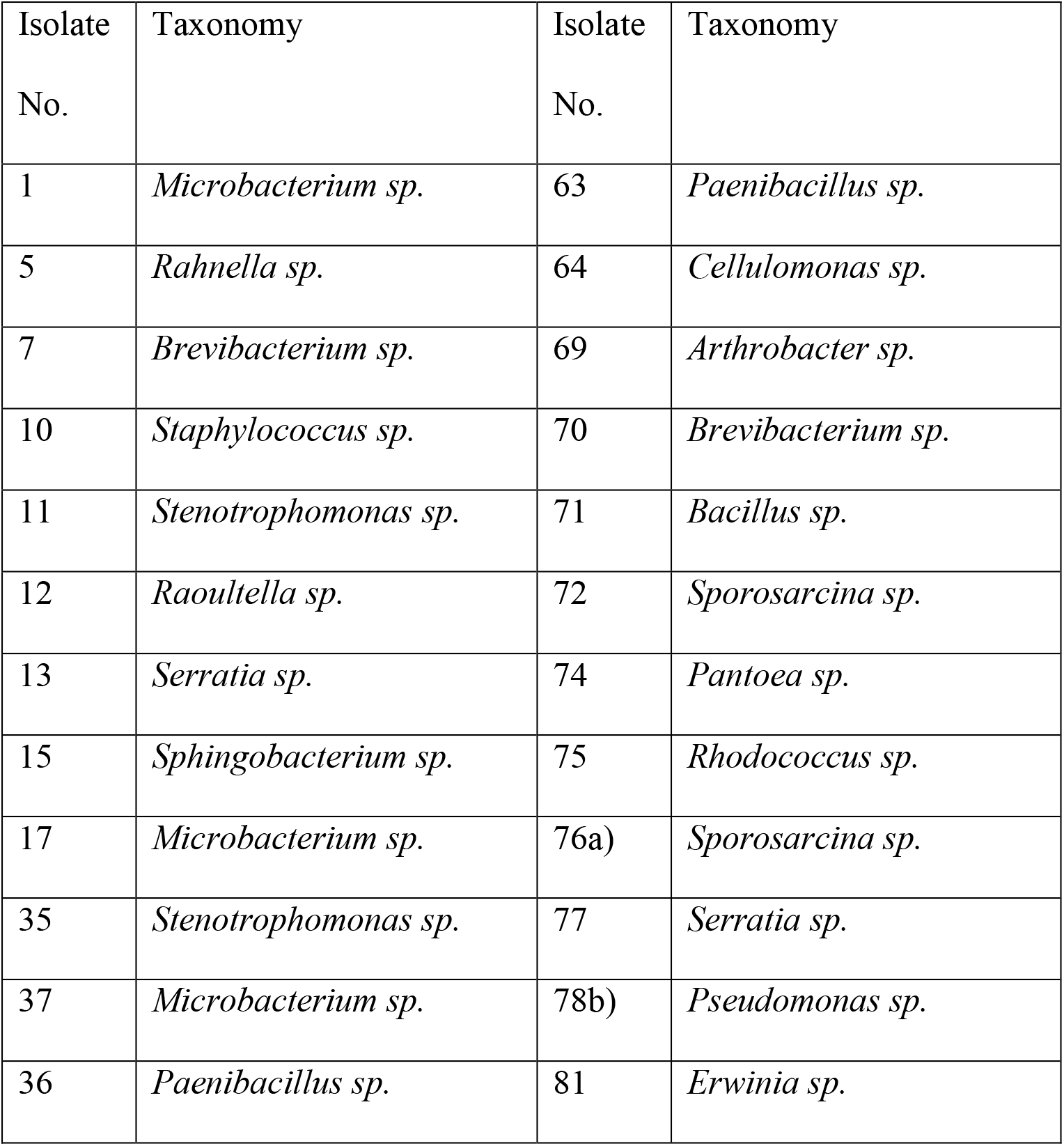
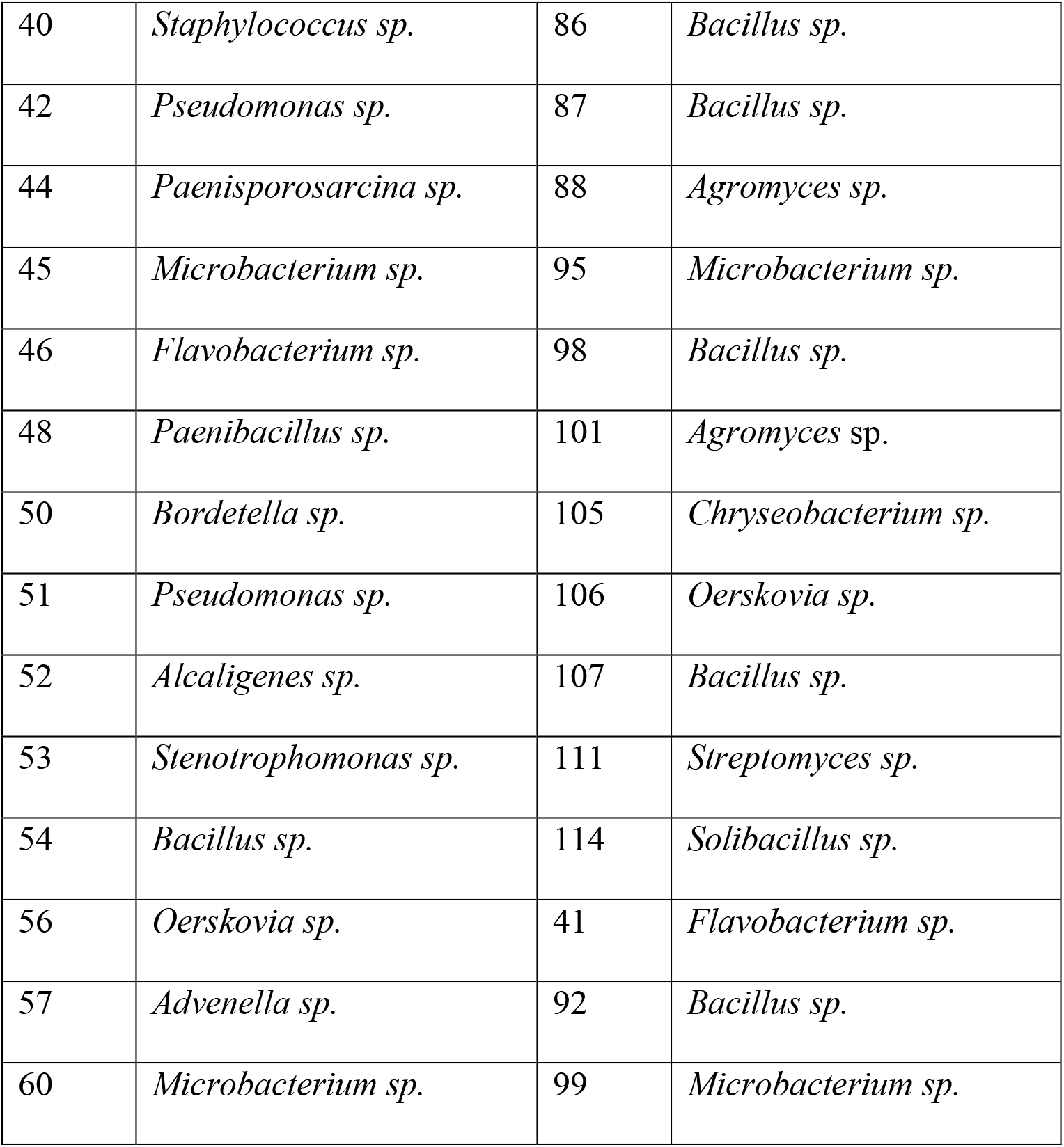
Bacterial isolates used for co-culturing experiments

| Isolate No. | Taxonomy | Isolate No. | Taxonomy |
| --- | --- | --- | --- |
| 1 | *Microbacterium sp.* | 63 | *Paenibacillus sp.* |
| 5 | *Rahnella sp.* | 64 | *Cellulomonas sp.* |
| 7 | *Brevibacterium sp.* | 69 | *Arthrobacter sp.* |
| 10 | *Staphylococcus sp.* | 70 | *Brevibacterium sp.* |
| 11 | *Stenotrophomonas sp.* | 71 | *Bacillus sp.* |
| 12 | *Raoultella sp.* | 72 | *Sporosarcina sp.* |
| 13 | *Serratia sp.* | 74 | *Pantoea sp.* |
| 15 | *Sphingobacterium sp.* | 75 | *Rhodococcus sp.* |
| 17 | *Microbacterium sp.* | 76a) | *Sporosarcina sp.* |
| 35 | *Stenotrophomonas sp.* | 77 | *Serratia sp.* |
| 37 | *Microbacterium sp.* | 78b) | *Pseudomonas sp.* |
| 36 | *Paenibacillus sp.* | 81 | *Erwinia sp.* |
| 40 | *Staphylococcus sp.* | 86 | *Bacillus sp.* |
| 42 | *Pseudomonas sp.* | 87 | *Bacillus sp.* |
| 44 | *Paenisporosarcina sp.* | 88 | *Agromyces sp.* |
| 45 | *Microbacterium sp.* | 95 | *Microbacterium sp.* |
| 46 | *Flavobacterium sp.* | 98 | *Bacillus sp.* |
| 48 | *Paenibacillus sp.* | 101 | *Agromyces sp.* |
| 50 | *Bordetella sp.* | 105 | *Chryseobacterium sp.* |
| 51 | *Pseudomonas sp.* | 106 | *Oerskovia sp.* |
| 52 | *Alcaligenes sp.* | 107 | *Bacillus sp.* |
| 53 | *Stenotrophomonas sp.* | 111 | *Streptomyces sp.* |
| 54 | *Bacillus sp.* | 114 | *Solibacillus sp.* |
| 56 | *Oerskovia sp.* | 41 | *Flavobacterium sp.* |
| 57 | *Advenella sp.* | 92 | *Bacillus sp.* |
| 60 | *Microbacterium sp.* | 99 | *Microbacterium sp.* |

After one week of incubation, mycelium-bound pigments of *S. lacrymans* were visible in all co-cultivation experiments. After 31 days, secreted pigments had emerged in 42 of 49 interactions. No pigment was observed in co-cultivations with isolates no. 7 (*Brevibacterium* sp.), 15 (*Sphingobacterium* sp.), 48 (*Paenibacillus* sp.), 50 (*Bordetella* sp.), 53 (*Stenotrophomonas* sp.), 72 (*Sporosarcina* sp.) and 105 (*Chryseobacterium* sp.).

*S. lacrymans* was also able to inhibit the growth of some bacterial isolates. Isolates 69 (*Arthrobacter* sp.; Fig. 9), 77 (*Serratia* sp.), 78 b (*Pseudomonas* sp.), and 64 (*Cellulomonas* sp.) showed reduced growth in the presence of the fungus compared to the control experiment. Isolates 75 (*Rhodococcus* sp.) and 71 (*Bacillus* sp.) were moderately restricted in growth during the co-cultivation with *S. lacrymans*.

**Figure 9.**
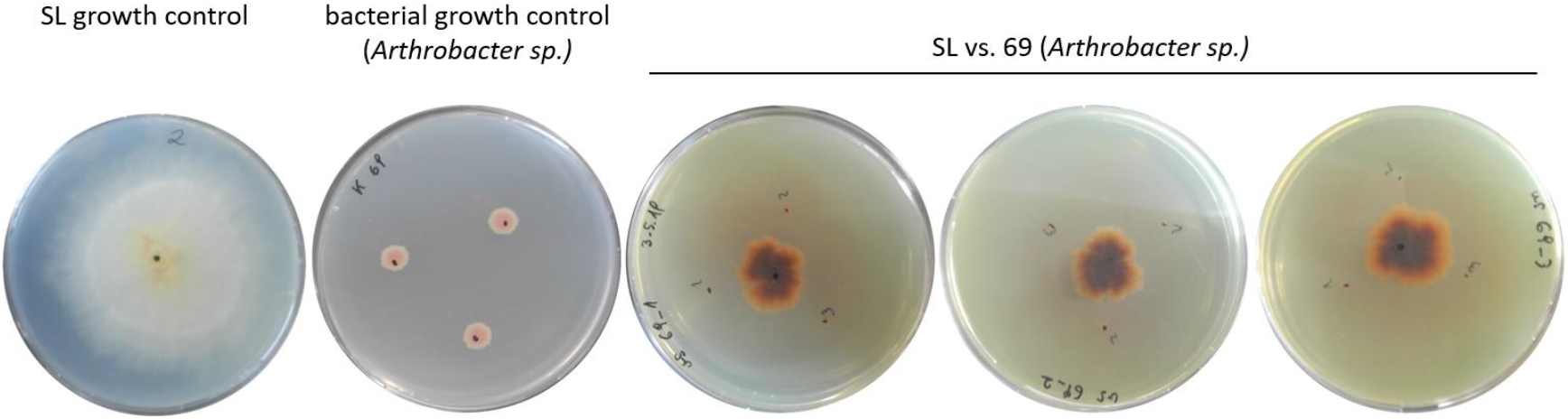
Inhibition of bacterial growth by *S. lacrymans*. Growth impairment of bacterial isolate 69 (*Arthrobacter* sp.) during co-cultivation with *S. lacrymans*. Pictures were taken after cultivation for 31 days on MM at 25 °C.

## Discussion

In the dead wood environment associated as well as co-occuring microorganisms originate from soil and/or air before they resettle on wood (2). In general, fungal fruiting bodies can harbor a broad spectrum of microorganisms including bacteria, yeasts, and filamentous fungi (41, 42). Previous studies based on cultivation dependent and independent methods revealed that the bacterial community in decaying wood is dominated by Proteobacteria, but found also bacteria belonging to the phyla Firmicutes, Actinobacteria, and Bacteroidetes (9, 43, 44). We provide evidence that the bacterial partners of *S. lacrymans* are dominated by Firmicutes, followed by Proteobacteria, Actinobacteria and Bacteroidetes. Moreover, different life stages of *S. lacrymans* (mycelia, rhizomorphs, basidiomata) were colonized by different bacterial communities. The bacterial community coexisting with *Hypholoma fasciculare* in decaying wood is dominated by Proteobacteria, followed by Acidobacteria, Firmicutes and Bacteroidetes (9). Co-existing bacteria of the mykorrhizal bitter bolete, *Tylopilus felleus*, were predominantly Proteobacteria and to a lesser extent Firmicutes, Bacteroidetes, Actinobacteria, and Cyanobacteria (30). These findings are similar to the findings of our expermiments. However, it can never be ecxluded that bacterial co-occurrence is independent of any causal relationships and therefore being the result of stochastic ‘mixing’ of *S. lacrymans* with bacteria. For instance exo-spore-forming bacteria such as *Streptomyces* sp. are known to be present in moisture-damaged building materials (45, 46). Such random co-occurances are making it challenging to distinguish true interactions from co-occurrence, especially as obviously bacteria growing on the substrate are isolated as well.

### Extension of isolation and cultivation experiments by FISH

In our study, we used a cultivation-based approach being well aware of the fact that most bacteria in environmental samples are uncultivable (47). To overcome potential drawbacks of this procedure, isolation and cultivation was complemented by FISH which allowed us to gain additional insights into the spatial distribution of bacteria on fungal tissues. We found evidence that environment-facing structures, like the hymenium of fruiting bodies, are primarily associated with bacteria while the number of bacteria decreases towards the inner parts (Fig. 4). This corresponds to our observations that hyphae inside the fruiting body appeared germfree and hence suitable for the isolation of *S. lacrymans* strains. The bacteria visualized with FISH were dominated by Firmicutes, while the abundance of β- and γ-Proteobacteria was lower (Fig. 6). This coincides well with the culture-dependent approach, where Firmicutes was also the most dominant phylum. Moreover, the number of CFU g^−1^ fruiting body tissue estimated based on the FISH results was comparable to the CFU counts. We analyzed two mature and one not yet fully developed fruiting body with FISH, and importantly, the total number of signals on the three fruiting bodies was similar, although the immature fruiting body showed a higher content of β- and γ-Proteobacteria. This indicates a development-stage-dependent shift of the bacterial composition on fruiting bodies, similar to what has previously been shown for *Cantharellus cibarius* (48). Our investigation of mycelial and rhizomorphic tissues of *S. lacrymans* with different single probes revealed individual bacterial cells but no spatial bacterial dispersal pattern (Fig. 5). We saw no hints for a particular dispersal pattern in comparison to the investigated fruiting bodies, and therefore excluded these tissue types from further multiplexing experiments. Bacteria were detected on the tissue surface as well as on hyphae and between fungal cells, but not within cells with wide lumen (e.g. rhizomorphs), as it is known for instance for *Mortierella elongata*, which harbors the endobacterium *Mycoavidus cysteinexigens* (49). Our analyses of *S. lacrymans* tissue revealed no embedded but individual bacterial cells, in contrast to the reindeer lichen *Cladonia arbuscula*, where bacterial cells were found embedded in a biofilm-like, continuous layer (50).

### Performance in enzymatic activity assays

Another benefit of isolation-based approaches is that the obtained bacterial isolates can be used for further experiments e.g. to assess their activities and influence on wood and timber degradation by *S. lacrymans*. As it is long known that bacteria cause structural changes in wood (51), we monitored the enzymatic repertoire of the obtained *S. lacrymans-*derived bacteria. Certain bacteria, particularly Actinobacteria (*Rhodococcus*), Firmicutes, Bacteroidetes (*Bacteroides*), Alpha-Beta- and Gammaproteobacteria (e.g. *Sphingomonas* and *Burkholdia*), have powerful cellulo-and pectinolytic enzyme systems as well as the capability to degrade lignocellulose, while pectinases play a key role in increasing wood permeability (51–53). We found that 10 out of 327 bacteria produced pectinase, amylase, cellulase, and xylanase including few isolates of *Serratia* sp., *Paenibacillus* sp., *Flavobacterium* sp., *Arthrobacter* sp., *Bacillus* sp., and *Pseudomonas* sp.. The enzymatic power of distinct bacteria such as *Serratia marcescens,* which is able to hydrolyse carboxymethylcellulose (54), *Bacillus polymyxa* that can hydrolyse pectin and holocellulose (55), and *Pseudomonas* sp. that are known for their phenol and aromatics degrading ability (56), is well documented. In addition, *Burkholderia* sp. are known as efficient mineral-weathering and nitrogen-fixing bacteria (57). It therefore might be worth to further characterize the microbial collection established in this study for the ability to fix nitrogen, as especially nitrogen is scarce in wood and fungi may overcome this deficiency by translocating the element from the surrounding environment where it is accumulated by N-fixing bacteria (58). Another interesting point would be to test the isolated bacteria for chitinolytic enzymes as this might give hints towards their ability to feed on fungal tissue.

### Single bacterial isolates caused growth inhibition of *S. lacrymans* and vice versa

Co-cultivations are frequently used to trigger production of bioactive metabolites by microorganisms, as axenic cultures do not reflect the natural situation (59). On the one hand bacteria may be antagonistic to fungi and competing for resources (23), on the other hand they can be beneficial, as certain bacterial metabolic products may function as growth factors for fungi (7) (Fig. S1). In our study, among the 50 tested bacterial isolates, 16 had no or a slightly positive effect on the mycelial growth of *S. lacrymans* in the co-cultivation assays, however this effect was very minute. Bacteria with the highest growth inhibiting effect on *S. lacrymans* were *Microbacterium* sp. (isolate no. 1, 45, and 95), *Arthrobacter* sp. isolate 69, *Bacillus* sp. (isolate no. 71 and 98), *Paenibacillus* sp. isolate 36, *Oerskovia* sp. isolate 106 and *Streptomyces* sp. isolate 95 (Fig. 8B; last nine isolates). Otherwise, *S. lacrymans* could inhibit the growth of some bacterial isolates too. We saw that genera, which cause growth impairment of *S. lacrymans,* and bacterial genera, which are decreased in their growth by the fungus, overlapped. This indicates that a cross-talk took place that resulted in the enhanced production of bioactive metabolites by the interaction partners. This over-production could impair the growth of the bacterial counterpart and, vice versa, the bacterium might start to produce substances that inhibit the fungus.

Another interesting observation was that the bacterial strains causing strong growth impairment of *S. lacrymans* also showed good performance in the enzyme activity test. This reveals that a broad repertoire of enzymatic features is useful in the warfare for habitat and nutrients. The fungal efforts in this warfare is for instance well noticeable by the selective effects of white-rot basidiomycetes on the bacterial community during wood-decay (23). Therefore, the competition between bacteria and fungi may influence the establishment and subsequent changes of the bacterial community.

## Conclusion

Our study aimed at a first characterization of mechanistic details of the *S. lacrymans* – bacteria interaction, and therefore cultures of the bacterial isolates were favored over mere DNA analysis, which simply gives information on the presence/absence of species. Importantly, both approaches, the culture-dependent assay and FISH microscopy, while based on different principles, gave similar results, and consequently, the outcome gains more meaningfulness. Nonetheless, future studies that include uncultured bacteria identified via metagenomic studies are needed, as well as studies on the fungal interaction partners of *S. lacrymans*. The exact mechanisms of the interaction of *S. lacrymans* with its bacterial microbiome remain also open. In summary, we provide the first insights of bacterial interaction partners of the dry-rot fungus *S. lacrymans* among which we found a dominance of Gram-positive bacteria. This could be emphasized by FISH experiments and the superiority of signals for the probe specific for Firmicutes. Bacterial isolates were tested for their enzymatic repertoire, especially for enzymes, which are relevant in the dead wood environment. The results of this experiments hint at the enormous power of bacteria in producing different enzyme types.

Especially single isolates of *Microbacterium* sp., *Flavobacterium* sp. *Arthrobacter* sp., and *Bacillus* sp. were, amongst others, promising candidates to further investigate their enzymatic repertoire and, based on their antagonistic activity, their secondary metabolites. Whether these bacteria have as well an effect on the wood-decaying properties of *S. lacrymans* remains to be revealed. This study provides new and relevant insights into the *S. lacrymans* bacterial microbiome and paves the way for future studies on recruitment, function and evolution of fungi-associated bacteria.

## Material and Methods

### Biological material

*Serpula lacrymans* fruiting bodies, mycelia and rhizomorphs were collected from infested timber in Austria, between summer 2018 and November 2019 (Tab. 1). These samples were used to isolate bacteria. To gain *S. lacrymans* pure cultures, a fruiting body from Innsbruck, collected in autumn 2018 (No. SLIBK2018), was cut with a sterile scalpel and material from the inside of the fruiting body was placed on malt extract agar (MEA, per l: 30 g malt extract, 3 g soy peptone, 15 g agar). Plates were incubated at 25 °C for 1 to 2 weeks.

### Isolation of bacteria from S. lacrymans fruiting bodies, mycelia and rhizomorphs

A total of 16 fruiting bodies, 9 rhizomorphs and 15 mycelia were collected in the period from summer 2018 to November 2019 (Tab. 1, Fig. 1A fruiting body, Fig. 1B mycelia, Fig. 1C rhizomorph). A defined amount (0.1 – 1 g) of fungal tissue was incubated for 10 minutes using a rotary shaker in 20 ml 0.9% sodium chloride – solution containing 0,015% Tween^®^ 80 (Sigma-Aldrich, St. Louis, United States of America) with 10 sterilized glass beads (Ø 4 mm). Dilution series up to 10^−6^ were prepared from this solution. 500 µl or 100 µl of each dilution step were plated on MEA plates, Tryptone soy agar (TSA, per l: 15 g casein peptone, 5 g soy peptone, 5g NaCl, 15 g agar), and R2A agar (from Roth, Karlsruhe, Germany; per l: 0.5 g yeast extract, 0.5 g peptone, 0.5 g casein hydrolysate, 0.5 g glucose, 0.5 g starch, 0.3 g potassium hydrogen phosphate, 0.04 g magnesium phosphate (MgSO_4_), 0.3 g sodium pyruvate, 15 g agar) and incubated at 25 °C for two to seven days. Plates were checked daily for growth of bacteria. We used a raster system to randomly isolate bacterial colonies for further analyses. All these pure cultures were dilution plated three times before further processing.

### Characterization of isolated bacteria

#### Molecular identification and phylogenetic relationship

Bacterial colony PCR was performed with the Red Taq 2x DNA Polymerase Master Mix (VWR, Radnor, USA; TRIS-HCl pH 8.5, (NH_4_)_2_SO_4_, 4 mM MgCl_2_, 0,2% Tween^®^ 20, each time 0.4 mM dNTPs, 0.2 units µl^−1^ Taq-Polymerase, inert color reagent). The PCR mixture contained 10 µmol of each of the primers 27F (5’ - AGA GTT TGA TCA TGG CTC A - 3’) and 1492R (1492R: 5’ - TAC GGT TAC CTT GTT ACG ACT T - 3’), 10.75 µl distilled water and 0.5 µl bovine serum albumin (BSA, 2%). Bacterial single colonies, not older than 3 days, were picked with a toothpick and added to the PCR-mixture. PCR conditions were 95 °C for 10 min, 30 cycles of 95 °C for 30 s, 53 °C for 30 s, 72 °C for 45 s, and a final incubation step at 72°C for 10 min. The PCR products were subject to agarose gel electrophoresis to confirm their correct size and subsequently sequenced using Microsynth sequencing service (Balgach, Switzerland).

Sequences were manually aligned using BioEdit Sequence Alignment Editor version 7.0.5.3 (60). Reference sequences were retrieved from the NCBI database. Phylogenetic trees were generated using PhyML (version 3.2, www.atgc-montpellier.fr/phyml) implemented Geneious version 9.1.8 with a general time reversible (GTR) substitution model and chi^2^ statistics to estimate branch support. Trees were optimized using the topology/length/rate function using NNI (next neighbor interchange) to search for the optimal tree topology. Sequences are available from the NCBI Sequence Read Archive with accession number MW089011 to MW089305 (for details see table S1 in the supplement).

Using an inverse morphotype approach, abundances of bacterial species (in CFU g^−1^) were estimated by counting single cells on agar plates of different dilution steps. Cells of equal morphological characteristics were assumed to belong to the same genera and CFUs were calculated by considering the dilution step. Only plates with bacterial clones identified by sequencing were included in the inverse morphotype assay. Different species displaying the same or a highly similar morphotype, could not be distinguished resulting in an approximation of the proportion of selected species within the individual samples but not “true” CFU counts of the individual morphotypes.

#### Biopolymer degradation tests

We tested our collection of bacterial isolates for different enzymatic activities that might be useful in the dead wood environment, like xylanases, cellulases, pectinases, and amylases. Bacteria that were able to depolymerize pectin, xylan, starch or cellulose showed a clear halo-like zone around their colonies.

#### Starch hydrolysis test

Starch agar (per l: 10 g potato starch, 5 g pancreatic digest of gelatin, 3 g beef extract, 15 g agar; pH 6.8 ±0.2) was used for detection of starch hydrolyzing microorganisms (61). Inoculated plates were incubated at 25 °C for 48 to 96 hours.

#### Pectinase activity test

Screening for pectinase activity was performed on pectinase screening agar (PSAM per l: 2 g NaNO_3_, 0.5 g KCl, 0.5 g MgSO_4_, 1 g K_2_HPO_4_, 0.5 g tryptone, 20 g agar, 10 g pectin; Tabassum (62). Inoculated plates were incubated at 25 ºC for 48 to 96 hours.

#### Cellulose degradation test

Cellulolytic activity was assessed on carboxymethyl cellulose agar (CMC agar; per l: 10 g peptone, 10 g carboxymethyl cellulose (Fluka), 2 g K_2_HPO_4_, 0.3 g MgSO_4_.7H_2_O, 2.5 g (NH_4_)_2_SO_4_, 2 g gelatin, 15 g agar; pH 6.8-7.2 (63)). Inoculated plates were incubated at 25 °C for 48 to 96 hours.

#### Xylan degradation test

Screening for xylanase activity was done using minimal agar medium (per l: 0.05 g MgSO_4_·7H_2_O, 0.05 g NaCl, 0.01 g CaCl_2_, 0.2 g yeast extract, 0.5 g peptone, 15 g agar (64) with 0.5% oat spelt xylan as the only carbon source (65). Inoculated plates were incubated at 25 ºC for 48 to 96 hours.

#### Enzyme Activity Evaluation

After incubation, all biopolymer-degradation test plates were flooded with ca. 10 ml 50 mM potassium iodide-iodine solution (Merck, Darmstadt, Germany) and incubated for 15 minutes at room temperature. Excess iodine solution was poured off and the plates were washed with 1 M NaCl (61). Another tested evaluation method was the flooding with 0.1% (w/v) congo red solution (Fluka, St. Louis; United States of America) followed by washing with 1 M NaCl after 15 minutes (66). Both evaluation methods worked equally well. The formation of a clear hydrolysis zone around bacterial colonies indicated enzymatic activity.

### Co-culture of *S. lacrymans* with isolated bacteria

Co-cultures were performed on 85 mm petri dishes containing 20 ml minimal media (MM) with a low nitrogen to carbon ratio (C:N = 400:1) similar to the nutritional value found in wood (67, 68). To prepare the bacterial inoculum for the co-cultivation experiments, bacterial strains were incubated at 25 °C for up to 7 days on MM before transferring them onto plates containing *S. lacrymans* strain (No. SLIBK2018). SLIBK2018 had been pre-grown for three weeks on MM agar plates at 25 °C, before transferring it to the center of the confrontation plate where it was allowed to grow for seven days at 25 °C before bacteria were added (Tab. 2). In total, 50 bacterial isolates across the taxonomic range were tested. Co-culture plates were incubated at 25 °C for 72 days. The first 14 days, plates were photographed and observed at least every second day, later once a week. The distance between the growing zone of *S. lacrymans* mycelia and bacterial colony was measured. Production of mycelium-bound and secreted pigments by the fungus were noted. We defined a bacterial strain as inhibitory on fungal growth when the colony size was less than 90% compared to the fungal growth control. Assignment to the category ‘strong inhibiting’ was done when growth was less than 60% compared to the control.

## Fluorescence In Situ Hybridization (FISH)

### Sampling and fixation conditions

*S. lacrymans* tissues (fruiting bodies, mycelia, and rhizomorphs) were washed with sterile distilled water and fixed with 4% Roti^®^-Histofix (Roth, Karlsruhe, Germany) for 1 hour. Samples were subsequently dehydrated for 1 hour with 50% ethanol at 4 °C and twice with 80% ethanol for 1 hour at 4 °C, followed by washing with ethanol abs. Material was stored at −20 °C until use.

#### Isolation of the surface community by cuticle tape lift

*S. lacrymans* mycelia and rhizomorphs were used for imprint preparations using double-sided adhesive tape (Tesafilm double sided 7.5 mm x 12 mm, Tesa SE, Norderstedt, Germany) following a modified protocol of Vorholt et al. (69). Double-sided tape was glued onto microscope slides, without touching the upper side of the tape. Fungal tissue was carefully flattened onto the sticky layer of the tape using an ethanol-wiped, clean glass stick (~ 1 mm diameter). Clean tweezers were used to remove the tissue. The tape imprints were fixed with 4% Roti^®^-Histofix (Roth, Karlsruhe, Germany) for 3 hours at room temperature. After incubation, the slides were dipped for 10 s into sterile double-distilled water and then subsequently dehydrated for 5 min each in 50%, 80% and 100% ethanol. After drying the slides with compressed air and an additional drying step for 5 min in the dark, the slides were stored at −20 °C.

#### FISH probes and fluorochromes

FITC-, 6FAM-, Cy3-, and Cy5-labelled probes were applied sequentially or simultaneously depending on the required formamide concentrations of the probe (Tab. 3). Samples stained with multiple probes (EUB338I-IIIw-CY3, BET42aw-6FAM, GAM42aw-6FAM and LGC354ABCw-CY5) were simultaneously analyzed, taking advantage of the non-overlapping emission wavelengths of the fluorochromes (max. excitation/emission in nm: 6FAM 492/518, FITC 490/525, Cy3 548/562 and Cy5 650/670).

**Table 3:**
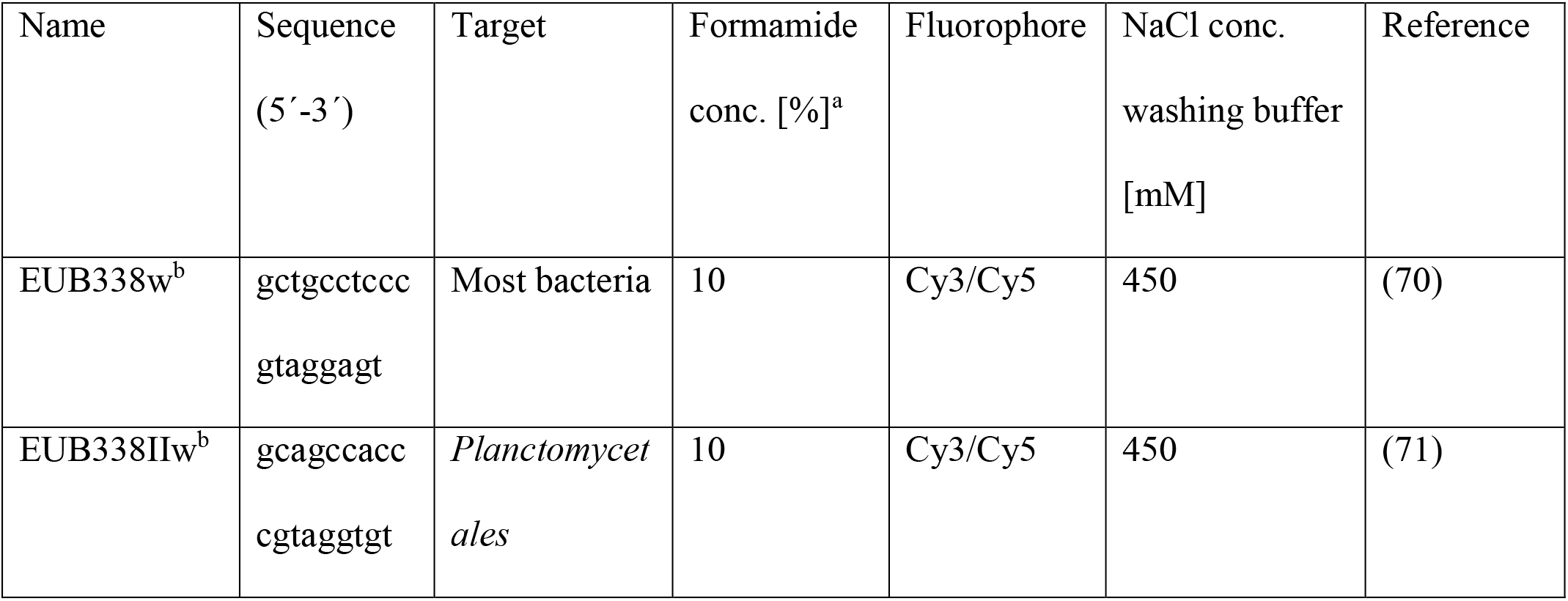
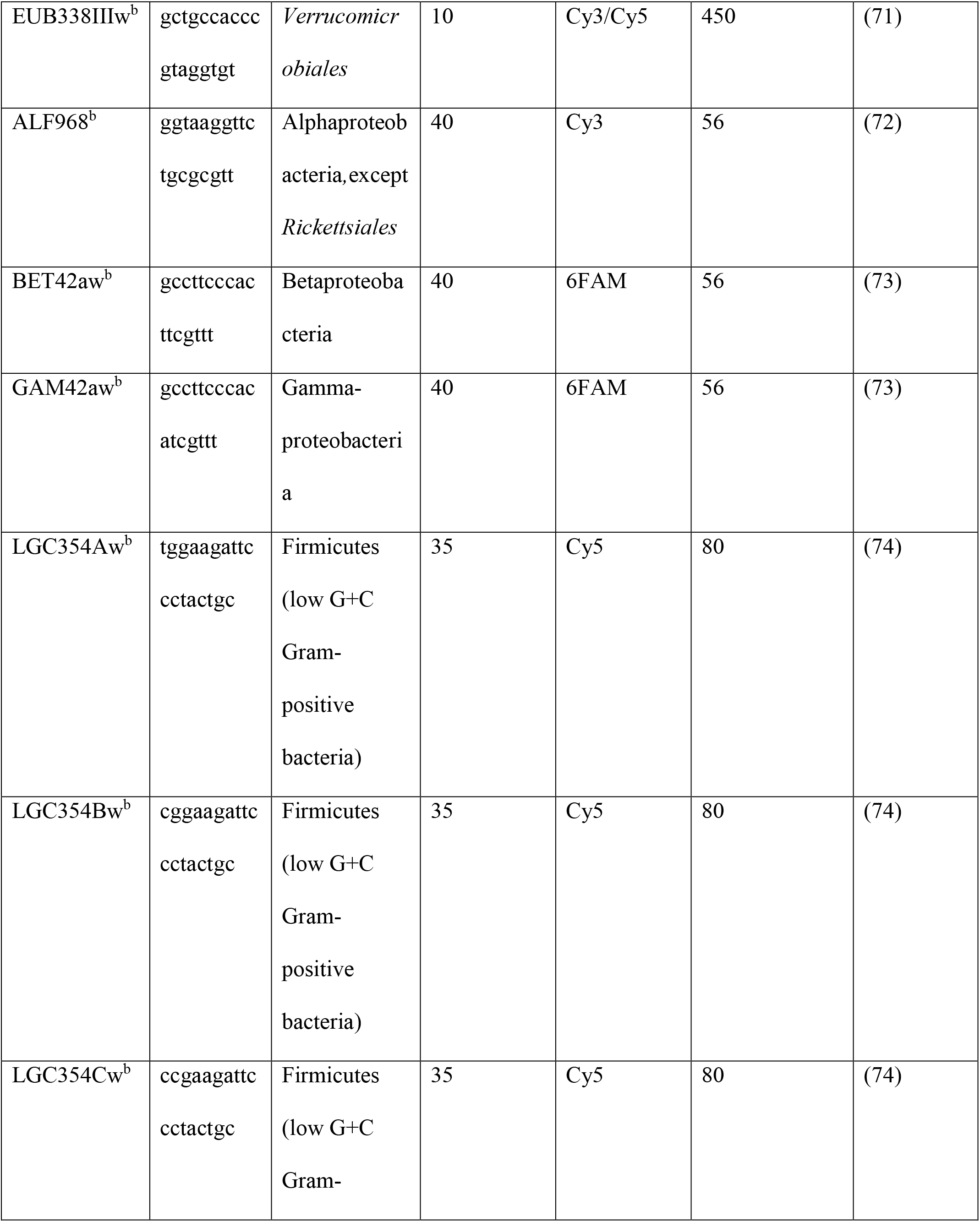
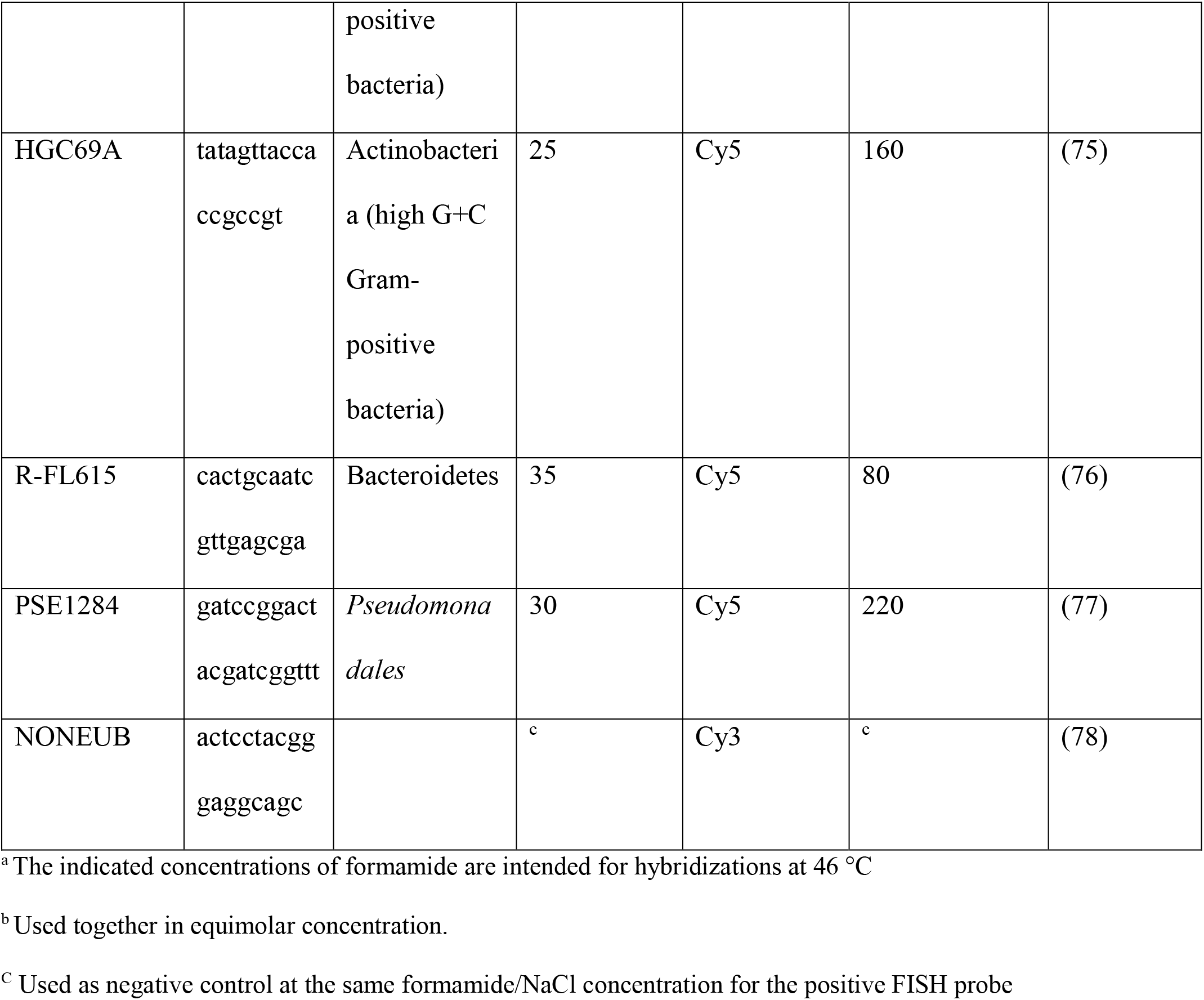
Characteristics of the probes used for Fluorescence in situ hybridization (FISH)

#### (In tube) FISH

Small pieces (10–15 mm length) of fixed and rehydrated tissue were embedded in O.C.T.TM Tissue-Teks (Sakura, Finetech Europe BV, Zoeterwoude, The Netherlands), rapidly frozen at −25 °C and cut into 30 µm-thick sections with a cryotome. The cryosections were placed in a 1.5 ml microcentrifuge tube and incubated with 1 mg ml^−1^ lysozyme (BioChemika, 105’000 U mg^−1^) for 20 min at room temperature to increase bacterial cell wall permeability for the FISH probes.

Additionally, a 10 mg ml^−1^ concentration of lysozyme was tested. The samples were then rinsed twice with ice-cold 1x PBS (8 g NaCl, 0.2 g KCl, 1.44 g Na_2_HPO_4_2H_2_O, 0.24 g KH_2_PO_4_ and 0.8 l distilled H_2_O). All hybridizations were performed at 46 °C overnight in a buffer containing 0.9 M NaCl, 0.02 M Tris-HCl (pH 8), 0.01% v/v sodium dodecyl sulphate (SDS) and 3 ng µl^−1^ of each probe. The concentration of ultrapure formamide (Invitrogen, Waltham, United States of America) for the FISH probes is given in Tab. 3. To completely immerse the investigated tissue type sections 60 µl of hybridization buffer was added. The hybridization buffer was removed the next day and the samples were washed twice with prewarmed (48 °C) washing buffer containing 0.02 M Tris-HCl (pH 8) and 30–450 mM NaCl (Tab. 3). The washing buffer was adjusted to 5 mM EDTA when the formamide concentration exceeded 20%. The sections were mounted onto a regular microscopic glass-slide.

#### Multiplex FISH

The presence of bacterial rRNA on *S. lacrymans* fruiting bodies was additionally investigated by sequential hybridization with probes specific for Eubacteria, EUB338 mix, Firmicutes (LGC354ABC mix), and β- and γ-Proteobacteria (BET42aw and GAM42aw) (Tab. 3).

Subsequent hybridizations were started with the probe with the highest formamide concentration, namely BET42aw and GAM42aw. Next, the probes specific for Firmicutes (LGC354A-C) were applied and finally the general bacterial probes were used (EUB338II-III). The incubation procedure was the same as previously described, with the exception that the incubation time at 46 °C was reduced to one hour and the probe hybridization was performed on teflon-precoated slides (increasement of the hybridization buffer volume to 240 µl).

#### FISH for imprints of the surface community on adhesive tape

Fixed and frozen double-side adhesive tape with imprints of fungal tissue was allowed to warm to room temperature in the dark to prevent precipitation of air humidity. Samples were once more dehydrated in an ethanol series as described above (section ‘Isolation of the surface community by cuticle tape lift’) to remove any residual water. Remaining ethanol was blown off using compressed air, and the slides were allowed to dry in the dark for 10 min. The tape was incubated with 1 mg ml^−1^ lysozyme (BioChemika, 105’000 U mg^−1^) for 20 min at room temperature to increase bacterial cell wall permeability for the FISH probes. For hybridization with probes specific for *Eubacteria* (EUB338 mix), the samples were subsequently overlaid with 240 µl hybridization buffer. The incubation procedure was the same as previously described, with the exception that the incubation time at 46 °C was reduced to three hours.

#### Nucleic acid and cell wall staining

FISH-stained sections and imprints on double-sided adhesive tape were incubated with 0.7 mg ml^−1^ 4,6-diamino-2-phenyl indole (DAPI) in the dark for 20 min for visualizing fungal nuclei at room temperature. After washing and drying, the sections were immediately mounted with VECTASHIELD^®^ (BIOZOL-Fit for Science, Eching, Germany) and finally observed under the fully automated Nikon inverted microscope Eclipse Ti2-E. The microscope was equipped with a Zyla sCMOS camera (Oxford Instruments, Abingdon, Great Britain), universal illumination system pE-4000 (CoolLED, Andover, Great Britain), intelligent polarizer Ti2-C-DICP-I (LWD 0.52), control unit TI2-CTRE, piezo nanopositioner combined with a Nano-Drive^®^ controller NIK-C2477 (Mad City Labs Inc., Madison WI, USA), motorized condenser turret TI2-C-TC-E, pillar for transmitted illumination TI2-D-PD, Epi-fluorescence module TI2-LA-FL and, control joystick TI2-S-JS), motorized DIC sextuple nosepiece with installed objectives (20x, 40x and 60x). Maximum projections of an appropriate number of about 0.5 mm-depth optical slices were applied to visualize fruiting body, mycelia and rhizomorphic sections (stacks) using standard settings. Fruiting bodies were cut into 30 µm thick slices and observed with 20x and 40x magnification, using the following filter sets: F66-413, F36-500, F36-720, F36-740 and F36-760. The digital images were further observed and analyzed with the NIS Elements Advanced Research Software from Nikon (Tokyo, Japan, version 5.11.01). Bacterial cells were counted manually after deconvolution (standard settings, type Landweber) using the Nikon software.

## Acknowledgments

We thank Nadine Präg for technical assistance with the molecular identification and Georg Walch for help with isolation of bacterial strains (both Department of Microbiology, University of Innsbruck). We also thank Willibald Salvenmoser and the Institute of Zoology (University Innsbruck) for usage of their Cryotome. This work was funded by the Leopold-Franzens-Universität Innsbruck and the doctoral programme ‘Biointeractions from basics to application’ (BioApp).

